# Cardiac events after macrolides or fluoroquinolones in patients hospitalized for community-acquired pneumonia: post-hoc analysis of a cluster-randomized trial

**DOI:** 10.1101/108407

**Authors:** D.F. Postma, C. Spitoni, C.H. van Werkhoven, L.J.R. van Elden, J.J. Oosterheert, M.J.M. Bonten

**Affiliations:** Julius Center for Health Sciences & Primary Care, University Medical Centre Utrecht, Heidelberglaan 100, 3508 GA Utrecht, the Netherlands; Department of Internal Medicine, Diakonessenhuis Utrecht, Bosboomstraat 1, 3582 KE Utrecht, the Netherlands; Department of Mathematics, Utrecht University, Budapestlaan 6, Room 601, 3584 CD Utrecht, the Netherlands; Department of Internal Medicine & Infectious Diseases, University Medical Centre Utrecht, Heidelberglaan 100, 3508 GA Utrecht, the Netherlands; Department of Pulmonary Medicine, Diakonessenhuis Utrecht, Bosboomstraat 1, 3582 KE Utrecht, the Netherlands; Department of Medical Microbiology, University Medical Centre Utrecht, Heidelberglaan 100, 3508 GA Utrecht, the Netherlands

**Author notes:** Corresponding author address: Julius Center for Health Sciences & Primary Care, University Medical Centre Utrecht, 3508 GA Utrecht, the Netherlands, |.

## Abstract

**Background:** Guidelines recommend macrolides and fluoroquinolones in patients hospitalized with community-acquired pneumonia (CAP), but their use has been associated with cardiac events.

**Objective:** To quantify associations between macrolide and fluoroquinolone use and cardiac events in patients hospitalized with CAP in non-ICU wards.

**Design:** Post-hoc analysis of a cluster-randomized trial

**Setting:** Six hospitals in the Netherlands

**Patients:** CAP patients admitted to non-ICU wards and without a cardiac event on admission

**Measurements:** Cause-specific hazard ratio’s (HR’s) were calculated for effects of time-dependent macrolide and fluoroquinolone exposure on cardiac events, defined as occurrence of new or worsening heart failure, arrhythmia, or myocardial ischemia during hospitalization.

**Results:** Cardiac events occurred in 146 (6.9%) of 2,107 patients and included episodes of heart failure (n=101, 4.8%), arrhythmia (n=53, 2.5%), and myocardial ischemia (n=14, 0.7%). Cardiac events occurred in 11 of 207 (5.3%), 18 of 250 (7.2%), and 31 of 277 (11.2%) patients exposed to azithromycin, clarithromycin, and erythromycin for at least one day, respectively, and in 9 of 234 (3.8%), 5 of 194 (2.6%), and 23 of 566 (4.1%) patients exposed to ciprofloxacin, levofloxacin, and moxifloxacin, respectively. Hazard ratios for any cardiac event, adjusted for confounding, were 0.89 (95% confidence interval (CI) 0.48 to 1.67), 1.06 (95% CI 0.61 to 1.83) and 1.68 (95% CI 1.07 to 2.62) for azithromycin, clarithromycin, and erythromycin, respectively, and adjusted hazard ratios were 0.86 (95% CI 0.47 to 1.57), 0.42 (95% CI 0.18 to 0.96) and 0.62 (95% CI 0.39 to 0.99) for ciprofloxacin, levofloxacin, and moxifloxacin, respectively. Erythromycin was associated with an adjusted hazard ratio of 2.08 (95% CI 1.25 to 3.46) for heart failure.

**Limitations:** Possibility of confounding by indication and observational bias

**Conclusions:** Among patients with CAP hospitalized to non-ICU wards, erythromycin use was associated with a 68% increased risk of hospital-acquired cardiac events, mainly heart failure. Levofloxacin and moxifloxacin were associated with a lower risk of heart failure.

*Registration:* The original trial was registered under ClinicalTrials.gov Identifier NCT01660204

*Funding Source:* The Netherlands Organization for Health Research and Development (ZONmw, Health care efficiency research, project id: 171202002).

## INTRODUCTION

The use of beta-lactam, macrolide, and fluoroquinolone antibiotics, either alone or in combination, is recommended in international guidelines for empirical treatment of patients hospitalized with community-acquired pneumonia (CAP).(1–3) However, the evidence-base for the addition of atypical coverage, especially macrolides, to beta-lactam antibiotics in patients admitted to non-ICU wards has been questioned.(4,5)

Results from observational studies have suggested that the use of macrolides (azithromycin, clarithromycin, and erythromycin) is associated with the occurrence of cardiovascular events, especially in patients with increased cardiovascular risk.(6–10) In a Danish study the relative risk for cardiac events after receiving azithromycin was not increased in the general population.(11) Still, the risk for myocardial infarction up to day 90 after admission increased (5.1% vs 4.4%; odds ratio 1.17 95% confidence interval (CI) 1.08 to 1.25) after azithromycin use in patients hospitalised with CAP. (7) Of note, azithromycin was associated with a lower risk of all-cause mortality. In another study, clarithromycin use during admission had an increased hazard ratio for cardiovascular events of 1.68 (95% CI 1.18 to 2.38) after one year follow-up among patients hospitalised with CAP.(8) Fluoroquinolones have been associated with an increased risk of arrhythmia in the general population, thus potentially increasing cardiovascular risk when treating CAP patients.(10,12)

Naturally, residual confounding cannot be excluded in these observational studies. However, the possibility of harm caused by macrolides or fluoroquinolones is likely to change the risk-benefit evaluation for atypical coverage in the empirical antibiotic treatment of CAP patients; particularly for macrolides in the context of limited evidence of benefit.

We recently demonstrated that a strategy of beta-lactam monotherapy was non-inferior to beta-lactams combined with a macrolide or fluoroquinolone monotherapy in terms of all-cause mortality for CAP patients admitted to non-intensive care unit (non-ICU) medical wards.(13) Here we present the results of a post-hoc analysis of these data on the association between specific macrolides and fluoroquinolones and the occurrence of cardiac events during hospitalization.

## METHODS

### Study setting

We used data from the *Community-Acquired Pneumonia: Study on the initial Treatment with Antibiotics of lower Respiratory Tract infections* (CAP-START) trial, a cluster-randomized cross-over trial comparing three empiric antibiotic strategies for the treatment of CAP. These strategies consisted of empirical treatment with beta-lactam monotherapy, beta-lactam/macrolide combination therapy, or fluoroquinolone monotherapy. The trial was performed in seven Dutch teaching hospitals in the Netherlands from February 2011 until October 2013, during which the strategies rotated in 4-month periods. In this study all patients who were admitted with a working diagnosis of CAP to a non-ICU ward for at least 24 hours were eligible for inclusion. Details about the methods and results of the trial have been previously published.(13,14)

For the current analysis we only included patients without evidence of a cardiac event at the time of hospital admission (i.e. in the emergency room).

Patients in the CAP-START trial gave written informed consent within 72 hours for data collection. Data collection for the current analysis was approved by the ethics review board of the University Medical Center Utrecht with a waiver for additional informed consent.

### Data collection

Demographic data, co-morbidities, clinical parameters, laboratory data, imaging results, and outcome data were collected from the medical records as part of the CAP-START trial by trained research nurses and were anonymously recorded in case record forms. During the course of the study, four investigators (DFP, CHvW: authors; LAM and KV: see acknowledgements) systematically collected additional data on the presence of cardiac and vascular comorbidities on admission, use of cardiovascular drugs, and the occurrence of cardiac events through chart review.

### Cardiac events

Pre-specified criteria for new or worsening cardiac events were used, as described previously.(15) We recorded three types of cardiac events: new or worsening arrhythmia, heart failure, and myocardial ischemia. New or worsening arrhythmia was defined as documentation in medical records or electrocardiogram (ECG) of newly recognized atrial fibrillation, flutter, supraventricular tachycardia, ventricular tachycardia or ventricular fibrillation. New or worsening heart failure was defined as clinical signs of new heart failure noted in medical records by treating physicians and a chest X-ray or CT-scan demonstrating evidence of heart failure. New or worsening myocardial ischemia was defined as documentation in medical records of at least two of three criteria: chest pain, acute electrocardiographic changes (ST segment and T wave changes without formation of Q waves, new Q waves or a clear loss of R waves), and/or elevated cardiac enzymes (CPK-MB, troponin-T). Cardiac events were considered worsening when therapeutic action (e.g. increase of diuretic dosage) was needed for already present cardiac disease.

First, admission and discharge letters were examined; if these did not contain detailed information, medical files were reviewed. In case of doubt, findings were presented in a plenary session to obtain group consensus (DFP, CHvW, and JJO: authors; LAM and KV: see acknowledgements) on the occurrence of an event.

### Antibiotic exposure

In the current analyses, macrolide and fluoroquinolone exposure were defined as time-varying exposures starting at the first day of antibiotic prescription until the end of admission. Azithromycin, clarithromycin, and erythromycin were the administered macrolides, and ciprofloxacin, ofloxacin, levofloxacin, and moxifloxacin were the administered fluoroquinolones. Ofloxacin was grouped with and referred to as ciprofloxacin because of the low number of ofloxacin users (n=1).

### Data analysis

We assessed the association between macrolide and fluoroquinolone exposure and cardiac events using an extended Cox proportional hazards model with time-varying covariates for exposure to these antibiotics. Antibiotic exposure was modelled as present from the starting date of the antibiotic prescription until the end of admission, as previously mentioned. Crude models included all six antibiotics (azithromycin, clarithromycin, erythromycin, ciprofloxacin, levofloxacin, and moxifloxacin) as time-dependent covariates and the different cardiac events (any cardiac event, heart failure, and arrhythmia) as outcomes; calculated hazard ratios are in comparison to patients who did not receive macrolide or fluoroquinolone antibiotics at any time during admission. Hospital discharge, transfer or death during admission led to right censoring in the analysis.

To calculate adjusted hazard ratios, we added the following confounders to our models: age, gender, a history of heart failure, number of other cardiac comorbidities, number of vascular comorbidities, hypertension, COPD, diabetes, a heart rate above 125bpm, hypotension, respiratory rate above 30bpm, serum sodium below 130mmol/L, arterial pH below 7.35, blood haematocrit below 30%, serum blood urea nitrogen above 11mg/dL, positive pneumococcal urinary antigen test, and serum C-reactive protein. The linearity of continuous variables was checked visually by plotting martingale residuals.

To assess the effect of the domain definition of clinical versus radiologically proven CAP, we repeated all analyses in the subgroup of patients with radiologically proven CAP.

Missing data were imputed by multiple imputation, except for data on respiratory rate, heart rate, and confusion at admission; the values for these variables were assumed to be normal when data were not documented in the medical records. P-values below .05 were considered statistically significant. All analyses were performed in R software, version 3.2.0.(16)

## RESULTS

### Patients

We included 2,107 patients who were admitted to non-ICU wards with a working diagnosis of CAP and without a cardiac event on admission (Figure 1). Baseline characteristics and outcomes, stratified by receipt of any macrolide or fluoroquinolone during admission, are displayed in Table 1. The median length of stay was 6 days (IQR 4-9 days). There were 146 (6.9%) patients with new cardiac events and 66 (3.1%) died during hospital stay (Table 1, Figure 1).

**Figure 1.**
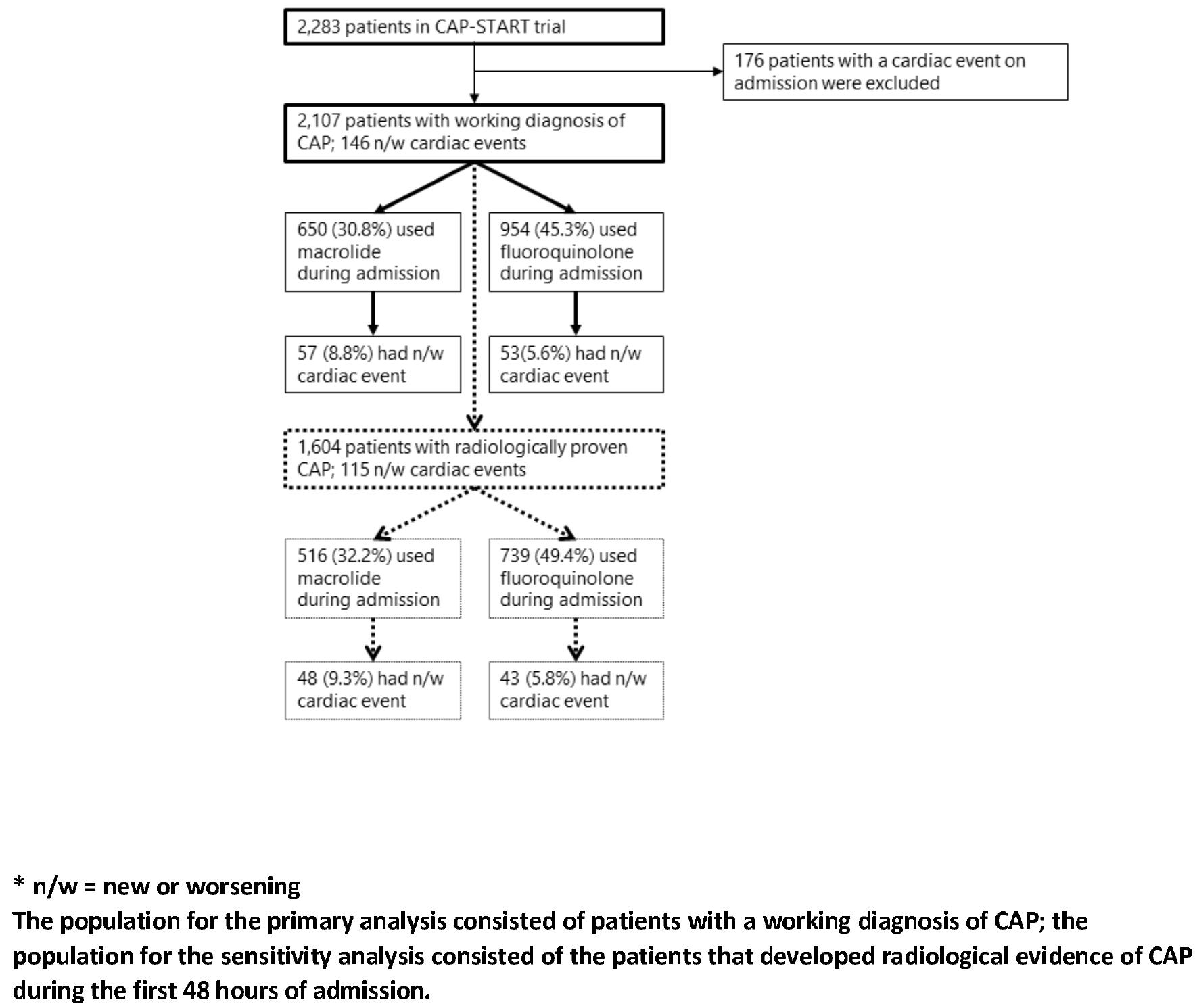
Inclusion of patients and cardiac event rates per population

**Table 1.** Baseline patient characteristics and outcomes stratified by use of any macrolide or fluoroquinolone during admission

|  | All patients | Use of any<br>macrolide<br>during admission | Use of any<br>fluoroquinolone<br>during admission | No<br>macrolide or<br>fluoroquinolone<br>during admission |
| --- | --- | --- | --- | --- |
| n | 2107 | 650 | 954 | 587 |
| Age | 69 (58;79) | 69 (57;78) | 69 (58;79) | 71 (60.5;79) |
| Male sex | 1217 (57.8%) | 388 (59.7%) | 556 (58.3%) | 332 (56.6%) |
| Nursing home residence<sup>1</sup> | 100 (4.8%) | 27 (4.2%) | 48 (5.1%) | 31 (5.4%) |
| Radiologically proven CAP | 1604 (76.1%) | 516 (79.4%) | 739 (77.5%) | 421 (71.7%) |
| **Co-morbidities** |  |  |  |  |
| History of cardiac disease | 733 (34.8%) | 193 (29.7%) | 342 (35.8%) | 227 (38.7%) |
| History of ischemic heart disease | 415 (19.7%) | 102 (15.7%) | 190 (19.9%) | 138 (23.5%) |
| History of atrial fibrillation | 299 (14.2%) | 88 (13.5%) | 141 (14.8%) | 86 (14.7%) |
| History of heart failure | 198 (9.4%) | 58 (8.9%) | 91 (9.5%) | 62 (10.6%) |
| History of vascular disease* | 458 (21.7%) | 130 (20.0%) | 219 (23.0%) | 133 (22.7%) |
| History of hypertension | 643 (30.5%) | 211 (32.5%) | 285 (29.9%) | 178 (30.3%) |
| History of COPD | 785 (37.3%) | 231 (35.5%) | 352 (36.9%) | 232 (39.5%) |
| History of diabetes | 339 (16.1%) | 83 (12.8%) | 175 (18.3%) | 97 (16.5%) |
| **Medication use<sup>2</sup>** |  |  |  |  |
| Use of antiplatelet agents | 575 (27.4%) | 186 (28.8%) | 252 (26.5%) | 170 (29.1%) |
| Use of anticoagulants | 345 (16.4%) | 85 (13.2%) | 165 (17.4%) | 104 (17.8%) |
| Use of antihypertensives | 1057 (50.4%) | 299 (46.3%) | 485 (51.1%) | 314 (53.7%) |
| Use of statins | 629 (30.0%) | 185 (28.6%) | 285 (30.0%) | 186 (31.8%) |
| **Severity scores** |  |  |  |  |
| PSI score ¶ | 83.7 ± 28.4 | 82.0 ± 27.9 | 84.6 ± 28.7 | 85.2 ± 28.4 |
| CURB65 score§ | 1 (1;2) | 1 (0;2) | 1 (1;2) | 1 (1;2) |
| **Outcomes** |  |  |  |  |
| n/w cardiac event | 146 (6.9%) | 57 (8.8%) | 53 (5.6%) | 42 (7.2%) |
| n/w arrhythmia | 53 (2.5%) | 19 (2.9%) | 21 (2.2%) | 14 (2.4%) |
| n/w heart failure | 101 (4.8%) | 46 (7.1%) | 34 (3.6%) | 27 (4.6%) |
| n/w myocardial ischemia | 14 (0.7%) | 2 (0.3%) | 7 (0.7%) | 6 (1.0%) |
| Transfer to other hospital | 12 (0.6%) | 1 (0.2%) | 7 (0.7%) | 5 (0.9%) |
| In-hospital mortality | 66 (3.1%) | 27 (4.2%) | 35 (3.7%) | 13 (2.2%) |

Values are medians (interquartile range) unless otherwise noted. Plus-minus values are means ± SD.
COPD denotes chronic obstructive pulmonary disease
\* includes cerebrovascular, peripheral artery, and thrombo-embolic disease.
¶ The PSI score uses 20 clinical measures to predict risk of death within 30 days, with results ranging from 0.1% (in patients with a score of 0–50) to 27.0% (in patients with a score >131).
§ The CURB-65 score is calculated by assigning 1 point each for confusion, uraemia (blood urea nitrogen ≥20 mg per decilitre), high respiratory rate (≥30 breaths per minute), low systolic blood pressure (<90 mm Hg) or diastolic blood pressure (≤60 mm Hg), and an age of 65 years or older, with a higher score indicating a higher risk of death within 30 days.
<sup>1</sup> Between 1-2.2% missing values for each group
<sup>2</sup> Between 0.3-0.6% missing values for each group

650 patients (30.8%) received at least one day of macrolide treatment, and 954 (45.3%) received at least one day of fluoroquinolones during admission. Treatment mostly started at the first day of admission (Table 2). Erythromycin was predominantly (94.9%) administered intravenously, whereas other macrolides were only used orally (as they are not available for intravenous administration in the Netherlands). The proportion of patients receiving fluoroquinolones that started with oral treatment on admission ranged from 30.4% for ciprofloxacin to 67.5% for moxifloxacin (Table 2). Patients receiving macrolides, as compared to the overall cohort, were less likely to have cardiac comorbidities (29.7% vs 34.8%), a history of diabetes (12.8% vs 16.1%), and anticoagulants (13.2% vs 16.4%) or antihypertensive medication (46.3% vs 50.4%). There were no obvious differences in baseline characteristics such as co-morbidities or use of cardiovascular medication for patients receiving fluoroquinolones as compared to the overall cohort. The disease severity upon admission, as represented by the PSI score or CURB-65 score, was comparable between patients with or without macrolides or fluoroquinolones (Table 1). The 146 patients with cardiac events developed 101 (4.8%) first episodes of heart failure, 53 (2.5%) of arrhythmia, and 14 (0.7%) of myocardial ischemia.

**Table 2.** Starting days and crude event rates for different macrolides and fluoroquinolones

| **Macrolides** |  | **Azithromycin** | **Clarithromycin** | **Erythromycin** |
| --- | --- | --- | --- | --- |
| Patients with antibiotic any time during admission |  | 207 | 250 | 277 |
| Starting day of antibiotic during admission ¶ |  | 1 (0-2) | 0 (0-1) | 0 (0-0) |
| Percentage starting antibiotic intravenously § |  | - | - | 263 (94.9%) |
| Cardiac event | a) Any type | 11 (5.3%) | 18 (7.2%) | 31 (11.2%) |
|  | b) Heart failure | 9 (4.3%) | 14 (5.6%) | 26 (9.4%) |
|  | c) Arrhythmia | 6 (2.9%) | 5 (2%) | 10 (3.6%) |
| **Fluoroquinolones** |  | **Ciprofloxacin** | **Levofloxacin** | **Moxifloxacin** |
| Patients with antibiotic any time during admission |  | 234* | 194 | 566 |
| Starting day of antibiotic during admission ¶ |  | 1 (0-3) | 0 (0-0) | 0 (0-0) |
| Percentage starting antibiotic intravenously |  | 76 (32.5%) | 111 (57.2%) | 394 (69.6%) |
| Cardiac event | a) Any type | 9 (3.8%) | 5 (2.6%) | 23 (4.1%) |
|  | b) Heart failure | 9 (3.8%) | 3 (1.5%) | 16 (2.8%) |
|  | c) Arrhythmia | 5 (2.1%) | 3 (1.5%) | 11 (1.9%) |

Values are numbers (percentages) unless otherwise noted. ¶ Median (interquartile range)
\* One patient received ofloxacin
§ Azithromycin and clarithromycin are not available for intravenous administration in the Netherlands

### Macrolide exposure and cardiac events

In the patients receiving macrolides (n=650), 207, 250, and 277 patients were exposed to azithromycin, clarithromycin, and erythromycin for at least one day, of which 11 (5.3%), 18 (7.2%), and 31 (11.2%) developed a cardiac event, respectively. The crude hazard ratio for a cardiac event after erythromycin use was 1.60 (95% CI 1.09; 2.36), and was 1.89 (95% CI 1.22; 2.91) for heart failure specifically. After adjustment for confounders hazard ratios were 1.68 (95% CI 1.07; 2.62) for any cardiac event and 2.08 (95% CI 1.25; 3.46) for heart failure. Adjusted hazard ratios for any cardiac event were 0.89 (95% CI 0.48; 1.67) and 1.06 (95% CI 0.61; 1.83) for azithromycin and clarithromycin, respectively. The numbers of cardiac events due to arrhythmia were too low for meaningful interpretations.(Table 3)

**Table 3.**
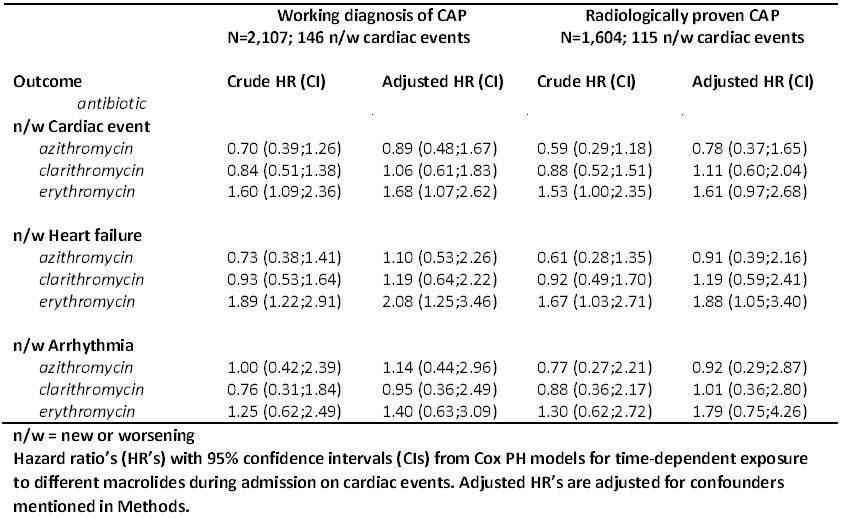
Hazard ratio’s for macrolides during admission

| Outcome | Working diagnosis of CAP<br>N=2,107; 146 n/w cardiac events |  | Radiologically proven CAP<br>N=1,604; 115 n/w cardiac events |  |
| --- | --- | --- | --- | --- |
|  | Crude HR (CI) | Adjusted HR (CI) | Crude HR (CI) | Adjusted HR (CI) |
| *antibiotic* |  |  |  |  |
| **n/w Cardiac event** |  |  |  |  |
| *azithromycin* | 0.70 (0.39;1.26) | 0.89 (0.48;1.67) | 0.59 (0.29;1.18) | 0.78 (0.37;1.65) |
| *clarithromycin* | 0.84 (0.51;1.38) | 1.06 (0.61;1.83) | 0.88 (0.52;1.51) | 1.11 (0.60;2.04) |
| *erythromycin* | 1.60 (1.09;2.36) | 1.68 (1.07;2.62) | 1.53 (1.00;2.35) | 1.61 (0.97;2.68) |
| **n/w Heart failure** |  |  |  |  |
| *azithromycin* | 0.73 (0.38;1.41) | 1.10 (0.53;2.26) | 0.61 (0.28;1.35) | 0.91 (0.39;2.16) |
| *clarithromycin* | 0.93 (0.53;1.64) | 1.19 (0.64;2.22) | 0.92 (0.49;1.70) | 1.19 (0.59;2.41) |
| *erythromycin* | 1.89 (1.22;2.91) | 2.08 (1.25;3.46) | 1.67 (1.03;2.71) | 1.88 (1.05;3.40) |
| **n/w Arrhythmia** |  |  |  |  |
| *azithromycin* | 1.00 (0.42;2.39) | 1.14 (0.44;2.96) | 0.77 (0.27;2.21) | 0.92 (0.29;2.87) |
| *clarithromycin* | 0.76 (0.31;1.84) | 0.95 (0.36;2.49) | 0.88 (0.36;2.17) | 1.01 (0.36;2.80) |
| *erythromycin* | 1.25 (0.62;2.49) | 1.40 (0.63;3.09) | 1.30 (0.62;2.72) | 1.79 (0.75;4.26) |

Risk estimates were similar in the subgroup analysis of patients with radiologically confirmed CAP.(Table 3) In most macrolide-exposed patients that developed cardiac events (n=45, 78.9%) macrolides were started on the day of admission. (Figure 2)

**Figure 2.**
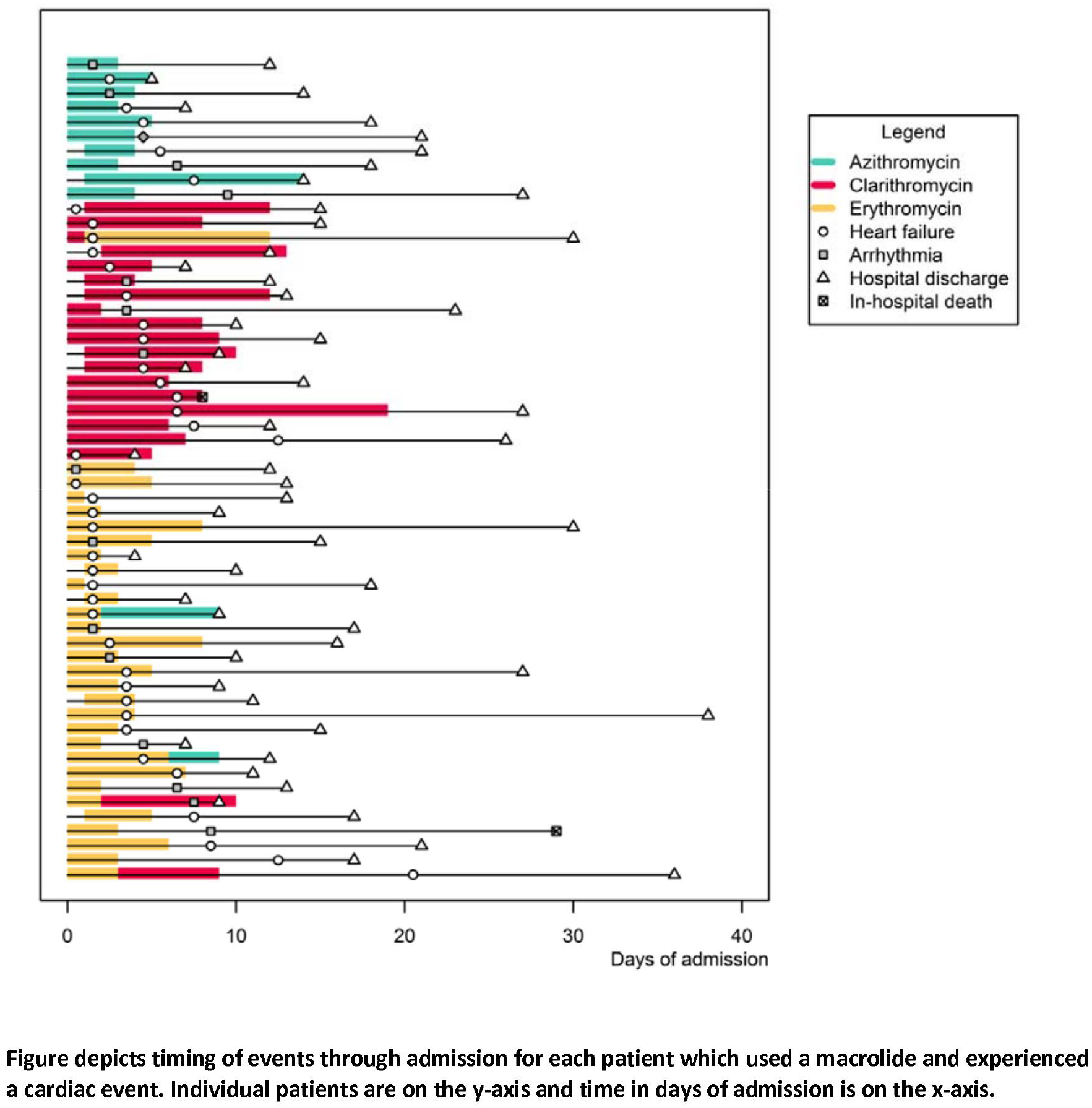
Timing of events in macrolide users with cardiac events

### Time-dependent exposure to fluoroquinolones

In the fluoroquinolone group (n=954), 234, 194, and 566 patients were exposed to ciprofloxacin, levofloxacin, and moxifloxacin, of which 9 (3.8%), 5 (2.6%), and 23 (4.1%) developed a cardiac event, respectively. (Table 2) Both levofloxacin and moxifloxacin were associated with lower risks of any cardiac event in crude analyses, with a hazard ratio of 0.40 (95% CI 0.18; 0.87) for levofloxacin and 0.56 (95% CI 0.36; 0.87) for moxifloxacin. Hazard ratios for heart failure specifically were 0.25 (95% CI 0.08; 0.80) for levofloxacin and 0.48 (95% CI 0.27; 0.84) for moxifloxacin. Associations for any cardiac event remained comparable after adjustment for confounders. The association between moxifloxacin and heart failure lost statistical significance after adjustment (Table 4). The numbers of cardiac events due to arrhythmia were too low for meaningful interpretations. Ciprofloxacin was not associated with a significantly changed hazard ratio for cardiac events.(Table 4)

**Table 4.** Hazard ratio’s for fluoroquinolones during admission

| Outcome | Working diagnosis of CAP<br>N=2,107; 146 n/w cardiac events |  | Radiologically proven CAP<br>N=1,604; 115 n/w cardiac events |  |
| --- | --- | --- | --- | --- |
|  | Crude HR (CI) | Adjusted HR (CI) | Crude HR (CI) | Adjusted HR (CI) |
| *antibiotic* |  |  |  |  |
| **n/w cardiac event** |  |  |  |  |
| *ciprofloxacin* | 0.77 (0.43;1.37) | 0.86 (0.47;1.57) | 0.81 (0.44;1.50) | 0.88 (0.46;1.67) |
| *levofloxacin* | 0.40 (0.18;0.87) | 0.42 (0.18;0.96) | 0.33 (0.12;0.91) | 0.29 (0.10;0.86) |
| *moxifloxacin* | 0.56 (0.36;0.87) | 0.62 (0.39;0.99) | 0.53 (0.32;0.86) | 0.63 (0.38;1.06) |
| **n/w Heart failure** |  |  |  |  |
| *ciprofloxacin* | 0.71 (0.36;1.44) | 0.85 (0.41;1.75) | 0.71 (0.34;1.50) | 0.81 (0.37;1.77) |
| *levofloxacin* | 0.25 (0.08;0.80) | 0.23 (0.07;0.76) | 0.12 (0.02;0.85) | 0.09 (0.01;0.66) |
| *moxifloxacin* | 0.48 (0.27;0.84) | 0.59 (0.33;1.06) | 0.43 (0.23;0.81) | 0.60 (0.32;1.16) |
| **n/w Arrhythmia** |  |  |  |  |
| *ciprofloxacin* | 0.83 (0.32;2.12) | 0.85 (0.32;2.26) | 0.73 (0.25;2.09) | 0.74 (0.25;2.21) |
| *levofloxacin* | 0.50 (0.15;1.64) | 0.57 (0.15;2.14) | 0.71 (0.21;2.39) | 0.72 (0.18;2.84) |
| *moxifloxacin* | 0.66 (0.33;1.34) | 0.68 (0.33;1.41) | 0.69 (0.32;1.46) | 0.73 (0.33;1.61) |

## DISCUSSION

In this study of 2,107 patients hospitalized with CAP to non-ICU wards, of which 146 (6.9%) developed a cardiac event during admission, erythromycin use was associated with a 68% higher hazard of cardiac events during hospital admission. This effect almost completely resulted from episodes of new or worsening heart failure. In contrast, levofloxacin and moxifloxacin were associated with a lower risk of cardiac events, mainly because of a lower risk of heart failure during admission.

This was a post-hoc analysis of a cluster-randomized cross-over study evaluating three antibiotic treatment strategies for patients hospitalized with CAP.(13) Baseline differences between patients who did or did not receive macrolides or fluoroquinolones during admission were small. If anything, classical cardiovascular risk factors were less prevalent in patients that received macrolides such as erythromycin. Hence, it is possible that physicians considered these antibiotics as potential risks for cardiac events in CAP patients, which may have attenuated the crude effect estimates in this analysis.(17) Consequently, the effect estimate between macrolides and cardiac events increased after adjustment for potential confounders. The findings were largely unchanged in patients with radiologically confirmed CAP. As associations between erythromycin use and competing outcomes such as in-hospital death or hospital discharge could change the interpretation of effect estimates, we also performed a competing risk analysis which did not change our results (data not shown; available with authors at request).(18)

Several biological mechanisms could explain the observed associations between erythromycin, levofloxacin, and moxifloxacin use and cardiac events, which could be attributed almost completely to occurrences of heart failure. The proposed explanation for this is the volume and sodium load associated with intravenous administration of erythromycin, compared to regimens including either other macrolides (that could only be used orally) or monotherapy with levofloxacin or moxifloxacin. The latter fluoroquinolones are also frequently given as oral treatment as the bioavailability of these antibiotics is known to be high.(21) Furthermore, erythromycin is a pro-arrhythmic drug, metabolized through CYP3A4 thus frequently interacting with other drugs, which can prolong the QT-interval.(19,20) Such effects would most likely occur shortly after start of treatment, and can lead to anything from self-limiting episodes of arrhythmia to severe arrhythmia or Torsade des Pointes, and consequently to demand ischemia and/or decompensated heart failure. Yet, the current analysis might have been underpowered to identify associations between use of any macrolide and occurrence of arrhythmias.

Our findings do not support previous findings of increased risks for cardiac events when using azithromycin or clarithromycin in hospitalized CAP patients.(7,8) In these studies all hospitalized patients with CAP, including those needing ICU-admission, were included with considerable longer follow-up periods for cardiac events, ranging from 90 days after discharge to one year after admission. In both studies, cardiac events mainly involved myocardial ischemia, which was a relatively infrequent event during hospitalization in our study. Moreover, fluoroquinolones have been associated with an increased risk for arrhythmia in the general population,(10,12) as mentioned before, our study was probably underpowered to properly assess this association.

Our post-hoc analysis has limitations. The occurrence of cardiac events during the course of the study was based on medical chart review in an open-label trial. This methodology has a risk for misclassification and observational bias. Random misclassification might have led to general underreporting of cardiac events, which would affect the precision of the effect estimates. More importantly, observational bias might have occurred if treating physicians would be more alert to report cardiac events in medical charts of erythromycin users, which would have biased the results from null. As this could also apply to registration of cardiac events by researchers, we used pre-specified criteria for recording cardiac events to reduce this bias. Another limitation could be residual confounding by indication. Although patient groups were comparable in terms of disease severity at baseline (Table 1), clinical stability during the first two to four days of admission is an important prognostic factor for adverse outcomes in CAP.(22,23) Therefore, clinical deterioration during the first days of admission might have stimulated treating physicians to add macrolides or fluoroquinolones, creating confounding by indication. Since changes in disease severity during these first days of admission were not determined, this could not be accounted for. Yet, almost all erythromycin users with a cardiac event (n=25, 80.6%) started with erythromycin on the day of admission.(Figure 2) Lastly, we did not have data on concomitant use of drugs interacting with macrolides or fluoroquinolones, nor did we determine specific causes of death. The latter could have increased the efficacy of end-point detection, and might have allowed discrimination between sudden cardiac death, cardiovascular death, and death due to other causes, although cause-of-death statements are not always interpreted unequivocally.(24)

### Conclusion

Our findings add to accumulating evidence that intravenous erythromycin use, but not oral azithromycin or clarithromycin use, increases the risk for cardiac events, especially heart failure, in patients hospitalized with CAP to non-ICU wards. Volume overload associated with intravenous administration of erythromycin in elderly is the most likely mechanism for heart failure. Inversely, this might explain why levofloxacin and moxifloxacin were associated with a lower risk of cardiac events, mainly heart failure, during admission. Although our study does not fully exclude confounding bias, findings remained largely unchanged in crude, adjusted, and sensitivity analyses. Together with the absence of survival benefit these findings should caution the use of intravenous erythromycin as empirical treatment of CAP patients admitted to non-ICU wards.

## ACKNOWLEDGEMENTS

We would like to thank all members of the CAP-START study group for securing the conduct of the study. Additional thanks go to L.A. Michielsen and K. Verbon for their help in data collection and preparation.

## Transparency declaration

The lead author affirms that this manuscript is an honest, accurate, and transparent account of the study being reported; that no important aspects of the study have been omitted; and that any discrepancies from the study as planned (and, if relevant, registered) have been explained.

## Data sharing

An anonymised dataset and full statistical code are available at request from the corresponding author.

## Author contributions

DFP participated in the design and conduct of the CAP-START study, performed the analysis, and wrote the current manuscript

CS participated in the design of the current analysis and performed parts of the analysis, and participated in writing the manuscript

CHvW participated in the design and conduct of the CAP-START study, and revised the current manuscript

LJRvE participated in the conduct of the CAP-START study and revised the current manuscript

JJO participated in the design and supervision of the CAP-START study, and revised the current manuscript

MJMB participated in the design and supervision of the CAP-START study, and supervised writing of the current manuscript

## REFERENCES

1. Mandell L a, Wunderink RG, Anzueto A, Bartlett JG, Campbell GD, Dean NC, et al. Infectious Diseases Society of America/American Thoracic Society consensus guidelines on the management of community-acquired pneumonia in adults. Clin Infect Dis [Internet]. 2007;44 Suppl 2(Suppl 2):S27–72. Available from:http://www.ncbi.nlm.nih.gov/pubmed/17278083

2. Lim WS, Baudouin S V, George RC, Hill AT, Jamieson C, Le Jeune I, et al.BTS guidelines for the management of community acquired pneumonia in adults: update 2009. Thorax. 2009;64 Suppl 3:iii1–i55.

3. Wiersinga WJ, Bonten MJ, Boersma WG, Jonkers RE, Aleva RM, Kullberg BJ, et al.SWAB/NVALT (dutch working party on antibiotic policy and dutch association of chest physicians) guidelines on the management of community-acquired pneumonia in adults. Neth J Med [Internet]. Department of Internal Medicine, University of Amsterdam, Amsterdam, the Netherlands.; 2012 Mar;70(2):90–101. Available from: http://www.ncbi.nlm.nih.gov/pubmed/22418758

4. Oosterheert JJ, Bonten MJM, Hak E, Schneider MME, Hoepelman IM. How good is the evidence for the recommended empirical antimicrobial treatment of patients hospitalized because of community-acquired pneumonia? A systematic review. J Antimicrob Chemother [Internet]. 2003 Oct 1 [cited 2015 Nov 23];52(4):555–63. Available from: http://jac.oxfordjournals.org/content/52/4/555

5. File TM, Marrie TJ. Does empiric therapy for atypical pathogens improve outcomes for patients with CAP? Infect Dis Clin North Am[Internet]. 2013 Mar [cited 2015 Nov 24];27(1):99–114. Available from: http://www.sciencedirect.com/science/article/pii/S0891552012001213

6. Ray WA, Murray KT, Hall K, Arbogast PG, Stein CM. Azithromycin and the risk of cardiovascular death. N Engl J Med [Internet]. 2012;366(20):1881–90. Available from: http://www.pubmedcentral.nih.gov/articlerender.fcgi?artid=3374857&tool=pmcentrez&rendertype=abstract

7. Mortensen EM, Halm E a, Pugh MJ, Copeland L a, Metersky M, Fine MJ, et al.Association of azithromycin with mortality and cardiovascular events among older patients hospitalized with pneumonia. JAMA [Internet]. 2014;311(21):2199–208. Available from: http://www.ncbi.nlm.nih.gov/pubmed/24893087

8. Schembri S, Williamson PA, Short PM, Singanayagam A, Akram A, Taylor J, et al. Cardiovascular events after clarithromycin use in lower respiratory tract infections: analysis of two prospective cohort studies. BMJ [Internet]. 2013 Jan [cited 2015 Sep 14];346:f1235. Available from: http://www.ncbi.nlm.nih.gov/pubmed/23525864

9. Ray WA, Murray KT, Meredith S, Narasimhulu SS, Hall K, Stein CM. Oral erythromycin and the risk of sudden death from cardiac causes. N Engl J Med [Internet]. 2004 Sep 9 [cited 2015 Sep 14];351(11):1089–96. Available from: http://www.ncbi.nlm.nih.gov/pubmed/15356306

10. Chou H-W, Wang J-L, Chang C-H, Lai C-L, Lai M-S, Chan KA. Risks of Cardiac Arrhythmia and Mortality Among Patients Using New-Generation Macrolides, Fluoroquinolones, and β-Lactam/β-Lactamase Inhibitors: A Taiwanese Nationwide Study. Clin Infect Dis [Internet]. 2015 Mar 15 [cited 2015 Sep 29];60(4):566–77. Available from: http://cid.oxfordjournals.org/content/60/4/566.full?etoc

11. Svanström H, Pasternak B, Hviid A. Use of azithromycin and death from cardiovascular causes. N Engl J Med [Internet]. 2013;368(18):1704–12. Available from: http://www.ncbi.nlm.nih.gov/pubmed/23635050

12. Lapi F, Wilchesky M, Kezouh A, Benisty JI, Ernst P, Suissa S. Fluoroquinolones and the risk of serious arrhythmia: a population-based study. Clin Infect Dis [Internet]. 2012 Dec [cited 2015 Sep 29];55(11):1457–65. Available from: http://www.ncbi.nlm.nih.gov/pubmed/22865870

13. Postma DF, van Werkhoven CH, van Elden LJR, Thijsen SFT, Hoepelman AIM, Kluytmans JAJW, et al. Antibiotic Treatment Strategies for Community-Acquired Pneumonia in Adults. N Engl J Med [Internet]. 2015 Apr 2 [cited 2015 Apr 2];372(14):1312–23. Available from: http://www.ncbi.nlm.nih.gov/pubmed/25830421

14. van Werkhoven CH, Postma DF, Oosterheert JJ, Bonten MJM. Antibiotic treatment of moderate-severe community-acquired pneumonia: design and rationale of a multicentre cluster-randomised cross-over trial. Neth J Med [Internet]. 2014 Apr [cited 2015 Mar 13];72(3):170–8. Available from: http://www.ncbi.nlm.nih.gov/pubmed/24846935

15. Corrales-Medina VF, Musher DM, Wells GA, Chirinos JA, Chen L, Fine MJ. Cardiac complications in patients with community-acquired pneumonia: incidence, timing, risk factors, and association with short-term mortality. Circulation [Internet]. 2012 Feb 14 [cited 2015 Mar 1];125(6):773–81. Available from: http://www.ncbi.nlm.nih.gov/pubmed/22219349

16. Team RDC. R: A Language and Environment for Statistical Computing [Internet]. R Foundation for Statistical Computing. 2011. Available from: http://www.r-project.org

17. Abo-Salem E, Fowler JC, Attari M, Cox CD, Perez-Verdia A, Panikkath R, et al. Antibiotic-induced cardiac arrhythmias. Cardiovasc Ther [Internet]. 2014 Mar [cited 2015 Sep 30];32(1):19–25. Available from: http://www.ncbi.nlm.nih.gov/pubmed/24428853

18. Andersen PK, Geskus RB, De witte T, Putter H. Competing risks in epidemiology: Possibilities and pitfalls. Int J Epidemiol. 2012;41(3):861–70.

19. Goldstein EJC, Owens RC, Nolin TD. Antimicrobial-Associated QT, Interval Prolongation: Pointes of Interest. Clin Infect Dis [Internet]. 2006 Dec 15 [cited 2015 Sep 29];43(12):1603–11. Available from: http://www.ncbi.nlm.nih.gov/pubmed/17109296

20. Guo D, Cai Y, Chai D, Liang B, Bai N, Wang R. The cardiotoxicity of macrolides: a systematic review. Pharmazie [Internet]. 2010 Sep [cited 2015 Sep 29];65(9):631–40. Available from: http://www.ncbi.nlm.nih.gov/pubmed/21038838

21. Zhanel GG, Fontaine S, Adam H, Schurek K, Mayer M, Noreddin AM, et al. A Review of New Fluoroquinolones⍰: Focus on their Use in Respiratory Tract Infections. TreatRespirMed. Department of Medical Microbiology, Faculty of Medicine, University of Manitoba, Winnipeg, Manitoba, CanadaDepartment of Clinical Microbiology, Health Sciences Centre, Winnipeg, Manitoba, CanadaDepartment of Medicine, Health Sciences Centre, Winnipeg, Man; 2006;5(1176–3450 (Print)):437–65.

22. Halm EA, Fine MJ, Marrie TJ, Coley CM, Kapoor WN, Obrosky DS, et al. Time to clinical stability in patients hospitalized with community-acquired pneumonia: implications for practice guidelines. JAMA [Internet]. 1998 May 13 [cited 2015 Sep 29];279(18):1452–7. Available from: http://www.ncbi.nlm.nih.gov/pubmed/9600479

23. Takada K, Matsumoto S, Kojima E, Iwata S, Ninomiya K, Tanaka K, et al. Predictors and impact of time to clinical stability in community-acquired pneumococcal pneumonia. Respir Med [Internet]. 2014 May [cited 2015 Sep 29];108(5):806–12. Available from: http://www.ncbi.nlm.nih.gov/pubmed/24589380

24. Smith Sehdev AE, Hutchins GM. Problems with proper completion and accuracy of the cause-of-death statement. Arch Intern Med [Internet]. 2001 Jan 22 [cited 2015 Sep 29];161(2):277–84. Available from: http://www.ncbi.nlm.nih.gov/pubmed/11176744

